# Morphological Hallmarks of White Matter Astrocytes Across Development

**DOI:** 10.64898/2026.09.23.753801

**Authors:** Kathleen H.M. Holmes, Mumu Fang, Katherine T. Baldwin

**Affiliations:** Department of Cell Biology and Physiology, University of North Carolina at Chapel Hill, Chapel Hill, NC 27599; Neuroscience Center, University of North Carolina at Chapel Hill, Chapel Hill, NC 27599

**Keywords:** astrocyte, white matter, morphology, glia

## Abstract

While the mechanisms underlying protoplasmic, or gray matter (GM), astrocyte morphogenesis have been extensively studied, far less is known about the development of fibrous, or white matter (WM), astrocytes. In particular, no study has comprehensively characterized the morphological development of WM astrocytes. To address this gap, we used an intracortical adeno-associated virus injection technique to sparsely label individual astrocytes in the mouse corpus callosum for morphological analysis across postnatal development. We found that astrocytes in the corpus callosum follow a distinct trajectory of morphological development from their GM cousins. They establish their branching complexity in the first two postnatal weeks and significantly increase in volume into adulthood. We further show that tiling is conserved in WM astrocytes and that a subset of mature astrocytes in the corpus callosum lack endfeet. These findings provide the first developmental framework for the morphogenesis of callosal astrocytes and establish a foundation for future studies investigating the molecular mechanisms regulating the development of WM astrocytes.

## Introduction

Astrocytes are a heterogenous population of morphologically and functionally complex glial cells that play varied and critical roles in brain homeostasis, synapse development, and neurovascular coupling. The functional complexity of astrocytes relies on their morphological complexity, as their elaborately branched arbors enable functional interactions with the many cells and structures in the tissue microenvironment (Baldwin et al., 2024; Torres-Ceja & Olsen, 2022). Altered astrocyte morphology is a hallmark feature of numerous neurological disorders (Endo et al., 2022; Genin et al., 2026; Jorge & Bugiani, 2019; Soto et al., 2023; Testen et al., 2018), underscoring the importance of proper astrocyte morphogenesis for overall brain health.

Nearly all our knowledge of astrocyte morphological development thus far relies on studies of protoplasmic astrocytes in gray matter (GM) brain regions, such as the cortex and hippocampus. This growing body of literature has established that GM astrocytes in the rodent brain develop postnatally, with the bulk of morphogenesis occurring during the first three postnatal weeks, in concert with synaptogenesis (Baldwin et al., 2021; Stogsdill et al., 2017; Watanabe et al., 2023). Several studies have identified mechanisms that regulate the morphological development of GM astrocytes, revealing the important role of astrocyte-synapse interaction in driving astrocyte development. Both direct astrocyte-synapse contacts and neuronally secreted factors have been shown to promote astrocyte-synapse interactions, enhance astrocytic morphological complexity and establish regional astrocyte specificity (Cheng et al., 2023; Holt et al., 2019; Morel et al., 2014; Stogsdill et al., 2017; Stork et al., 2014; Takano et al., 2020; Tan et al., 2023; Xie et al., 2022). Strikingly, only two neuron-independent mechanism have been identified: gap junction-mediated interactions between cortical astrocytes (Baldwin et al., 2021) and astrocyte self-recognition (Lee et al., 2025) are critical for their morphological maturation. Collectively, these studies indicate that GM astrocytes and synapses engage in a continuous, reciprocal feedback loop that shapes their development and specialization.

The first descriptions of astrocyte morphology in the late 1800s noted two astrocyte subtypes based on morphological differences (Köhler et al., 2021): protoplasmic astrocytes which reside in the GM and are characterized by complex, bushy arbors, and fibrous astrocytes which reside in the white matter (WM) and are less complex, with long spindly processes. Additional morphologically distinct subtypes have been identified since, including Bergmann glia in the cerebellum and Müller glia in the retina. Notably, these morphologically distinct subtypes reside in distinct tissue microenvironments. GM is rich in neuronal cell bodies and synapses that mediate information processing, while white matter (WM) contains myelinated axon tracts that transmit information between brain regions. Accordingly, astrocytes in GM and WM are specialized to support these functions. The complex arbors of protoplasmic astrocytes allow for extensive bidirectional communication with synapses (Farizatto & Baldwin, 2023; Lawal et al., 2022). In contrast, fibrous astrocytes in WM are specialized to support myelination and axonal signal propagation. They form direct gap junction-mediated interactions with oligodendrocytes (Orthmann-Murphy et al., 2008; Tress et al., 2012), regulate oligodendrocyte maturation (Moore et al., 2011; Padovani-Claudio et al., 2006; Stankoff et al., 2002), and supply essential lipids to the myelin sheath (Camargo et al., 2017). At nodes of Ranvier, WM astrocytes buffer K+ (Kalsi et al., 2004) and actively participate in Ca²⁺-mediated signaling to modulate the speed of signal propagation along axons (Hamilton et al., 2008; Lezmy et al., 2021). In addition to these known functional differences, GM and WM astrocytes are transcriptionally distinct further highlighting their regional specialization (Bocchi et al., 2025; Hasel et al., 2025).

Despite the importance of WM astrocytes for WM function, there is no source detailing the morphological features of WM astrocytes across development and the mechanisms that regulate their morphogenesis are unknown. This knowledge gap is significant, as the human brain is approximately fifty percent WM by volume (Allen et al., 2003) and many brain disorders feature WM pathology (Fitzgerald et al., 2019; Jorge & Bugiani, 2019; Nasrabady et al., 2018; Ozarkar et al., 2024). To address this gap, we conducted an in-depth analysis of WM astrocyte morphology across postnatal mouse development, including gross astrocyte morphology, astrocyte branching architecture, endfeet, tiling, and astrocyte-axonal contact.

## Results

### WM astrocytes show protracted development, flattened shape, and reduced complexity compared to GM astrocytes

To assess WM astrocyte morphology in the developing mouse brain, we conducted a thorough analysis of individual astrocyte morphology across postnatal development in the corpus callosum, the large subcortical WM tract which connects the brain’s right and left hemisphere. Astrocytes densely carpet the brain making it challenging to evaluate individual cell morphology with labeling strategies that uniformly target all astrocytes. To label individual astrocytes for morphological analysis, we utilized an adeno-associated viral (AAV) labeling strategy as previously described (Eaker et al., 2026) to specifically and sparsely label astrocytes with membrane-targeted mCherry-CAAX throughout the postnatal mouse brain (Fig. 1A). We collected brains at four key timepoints for astrocyte and corpus callosum development: 1) P8, a pre-myelination timepoint (Ozarkar et al., 2024), 2) P14, the peak of callosal axon development (De León Reyes et al., 2019; Mizuno et al., 2007), 3) P21, morphological maturation of gray matter astrocytes in the cortex ((Baldwin et al., 2021; Stogsdill et al., 2017; Watanabe et al., 2023), and 4) P70, an adult tissue state featuring pruned axons and more established myelination (Baloch et al., 2009; De León Reyes et al., 2019; Sturrock, 1980). To compare the morphology and developmental trajectory of astrocytes in GM and WM, we acquired confocal images of whole individual astrocytes between layers 2 and 5 of the somatosensory cortex and the medial corpus callosum from the same tissue sections. We focused our studies on the somatosensory cortex as this cortical region sits above the thickest parts of the corpus callosum and on layers 2 through 5 to achieve a broad sampling of cortical GM astrocytes. We avoided layer 1 astrocytes close to the pial surface and layer 6 astrocytes close to the corpus callosum. We focused on the medial corpus callosum, the continuous tract beneath the motor cortexes and the primary somatosensory cortexes medial of the barrel cortex (Supplemental Fig. 1), to minimize potential impacts of corpus callosum thickness on astrocyte morphology.

**Figure 1:**
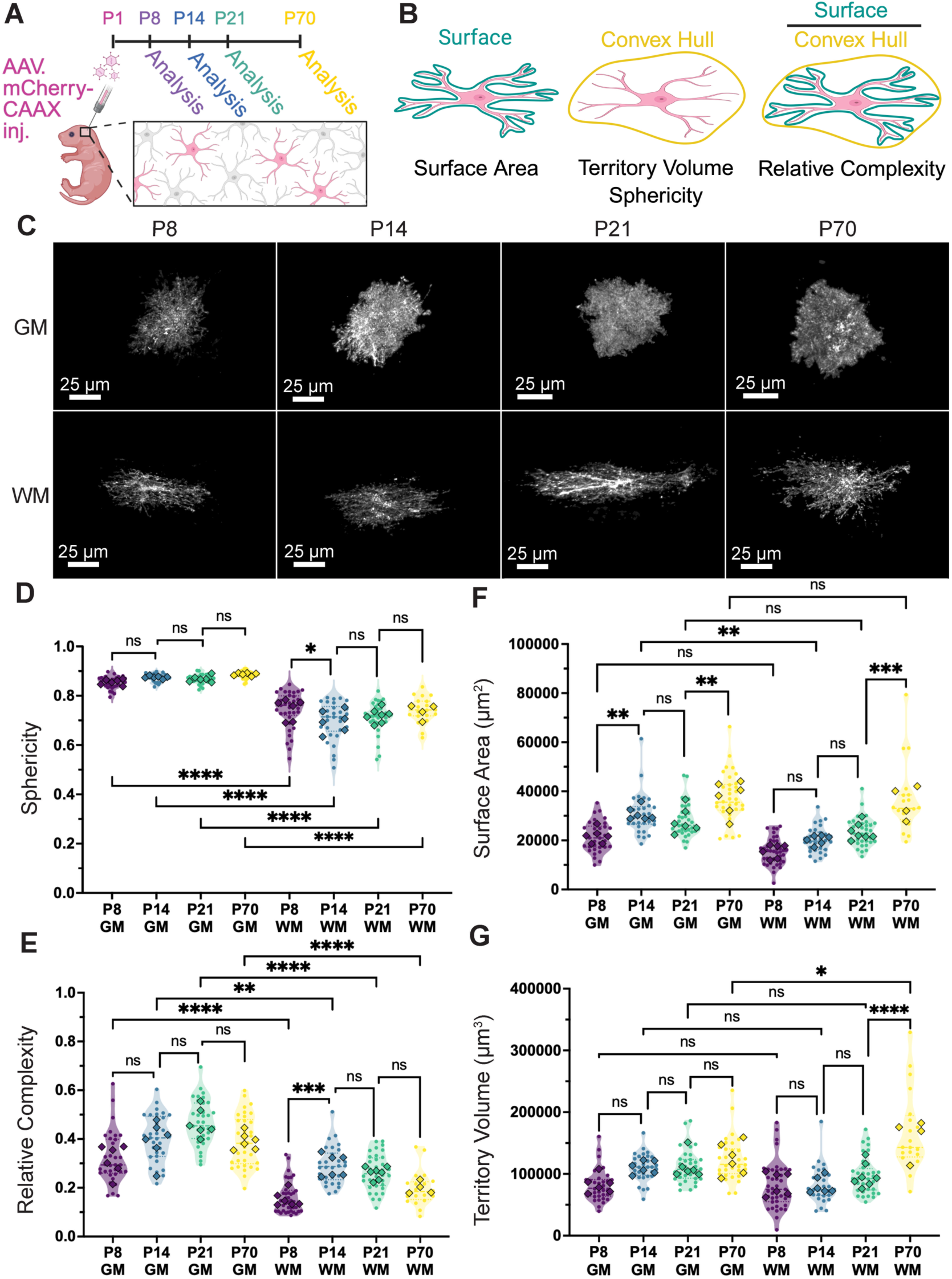
WM astrocytes show protracted development, flattened shape, and reduced complexity compared to GM astrocytes. **A)** Experimental overview showing AAV injection to generate brains with sparsely labeled astrocytes and collection timepoints. Created with BioRender.com. **B)** Visual descriptions of the elements in IMARIS used to developed four key morphology metrics: Surface Area, Territory Volume, Sphericity and Relative Complexity. Created using BioRender.com. **C)** Representative confocal images of whole, individual astrocytes expressing mCherry-CAAX in the cortex (GM) and corpus callosum (WM) at P8, P14, P21, and P70. The astrocyte channel was masked using the surface generated in IMARIS to focus on the analyzed astrocyte in each image. Scale bar, 25 µm. **D-F)** Violin plots showing 3D morphology analysis of astrocytes at P8, P14, P21 and P70 and in GM and WM. Data for **D)** sphericity **E)** relative complexity **F)** surface area and **G)** territory volume is presented with individual cells as dots and animal averages as diamonds. 3-9 cells per animal with 4-7 animals per condition. Nested one-way ANOVA, Sidak’s post-test. ns = p<0.05, * = p≤0.05, ** = p≤0.01, ***= p≤0.001, **** = p≤0.0001.

We used established assays (Eaker & Baldwin, 2022) to perform a multipoint assessment of astrocyte morphology, collecting measurements for territory volume, surface area, relative complexity, and sphericity (Fig. 1B) from GM and WM astrocytes at all time points. In contrast to the spherical, bushy arbors of GM astrocytes, WM astrocytes at all timepoints displayed a noticeably flattened shape and less-dense arbor (Fig. 1C). Accordingly, WM astrocyte sphericity (Fig. 1D) and relative complexity (Fig. 1E) were significantly lower than their GM counterparts across timepoints. Notably, we found no difference in maximum surface area between GM and WM astrocytes (Fig. 1F) suggesting astrocytes across tissues may reach an upper limit to the amount of membrane they can support. Within time course comparisons revealed that WM astrocytes experience a significant increase in relative complexity between P8 and P14 (Fig. 1E) and that their territory volume and surface area grows significantly between P21 and P70 (Fig. 1G). Previous studies that compared only two time points found significant increases in GM astrocyte territory volume and complexity between P7 and P21 (Stogsdill et al., 2017) with no change in territory volume between P21 and P70 (Baldwin et al., 2021). Our data are consistent with these findings, though the significance is masked upon correction for multiple comparisons. Uncorrected p-values (Supplemental table 1) show a significant increase in territory volume between P8 and P14 and P8 and P21, but no significant difference in GM territory volume between P14 and P21 or P21 and P70. These results demonstrate that GM astrocytes reach their full territory volume by the end of the second postnatal week, though complexity continues to increase through P21 and decreases slightly by P70 (Supplemental Table 1).

A comparison of GM and WM astrocyte developmental time courses reveals distinct developmental trajectories. GM astrocytes grow in volume and complexity simultaneously from P8 – P14 and continue to increase in complexity from P14 – P21 after reaching their final territory size (Fig. 1E, 1G). Conversely, WM astrocytes first display a sharp increase in complexity from P8-P14 (Fig. 1E), followed by a prolonged period of territory growth from P21 – P70 (Fig. 1G). While we did not observe any significant changes in WM astrocyte complexity during this time period, the relative complexity measurement used in this analysis does not directly measure branching architecture and, therefore, cannot capture any subtle changes in branching organization that may accompany the increase in volume.

### Branching complexity of WM astrocytes in unchanged between P21 and P70

To directly evaluate the branching complexity of WM astrocytes at P21 and P70, we conducted *in vivo* Sholl analysis of WM astrocytes at P21 and P70 in flat mounted corpora callosa. The sheet-like structure of the corpus callosum and flattened shape of WM astrocytes places the largest face of the astrocyte in the x-z plane for both coronal and sagittal sections, typically used for analysis of GM astrocytes. The resolution along the z-axis (∼0.5 µm) is lower than the x and y axes (∼0.25 µm), resulting in a “stretching” of 3D-rendered cell volumes in the z-dimension. To analyze WM astrocyte branching complexity with the largest face of the astrocyte in the x-y plane, we developed a protocol to microdissect the corpus callosum from fixed brains with AAV mCherry-CAAX injection (Fig. 2A). We then performed immunolabeling and flat mounted the tissue onto the slide such that the rostral-caudal and medial-lateral axes, which are the largest dimensions of astrocytes in the corpus callosum, lay in the x-y plane of the image (Fig. 2B). We collected high-resolution confocal images of mCherry-CAAX-expressing astrocytes (Fig. 2C) and adapted our previously described trained machine learning model in Imaris (Coble et al., 2026) to construct 3D filaments of WM astrocytes for subsequent 3D Sholl analysis (Gastinger, Oxford Instruments, 2021;Tan et al., 2023).

**Figure 2:**
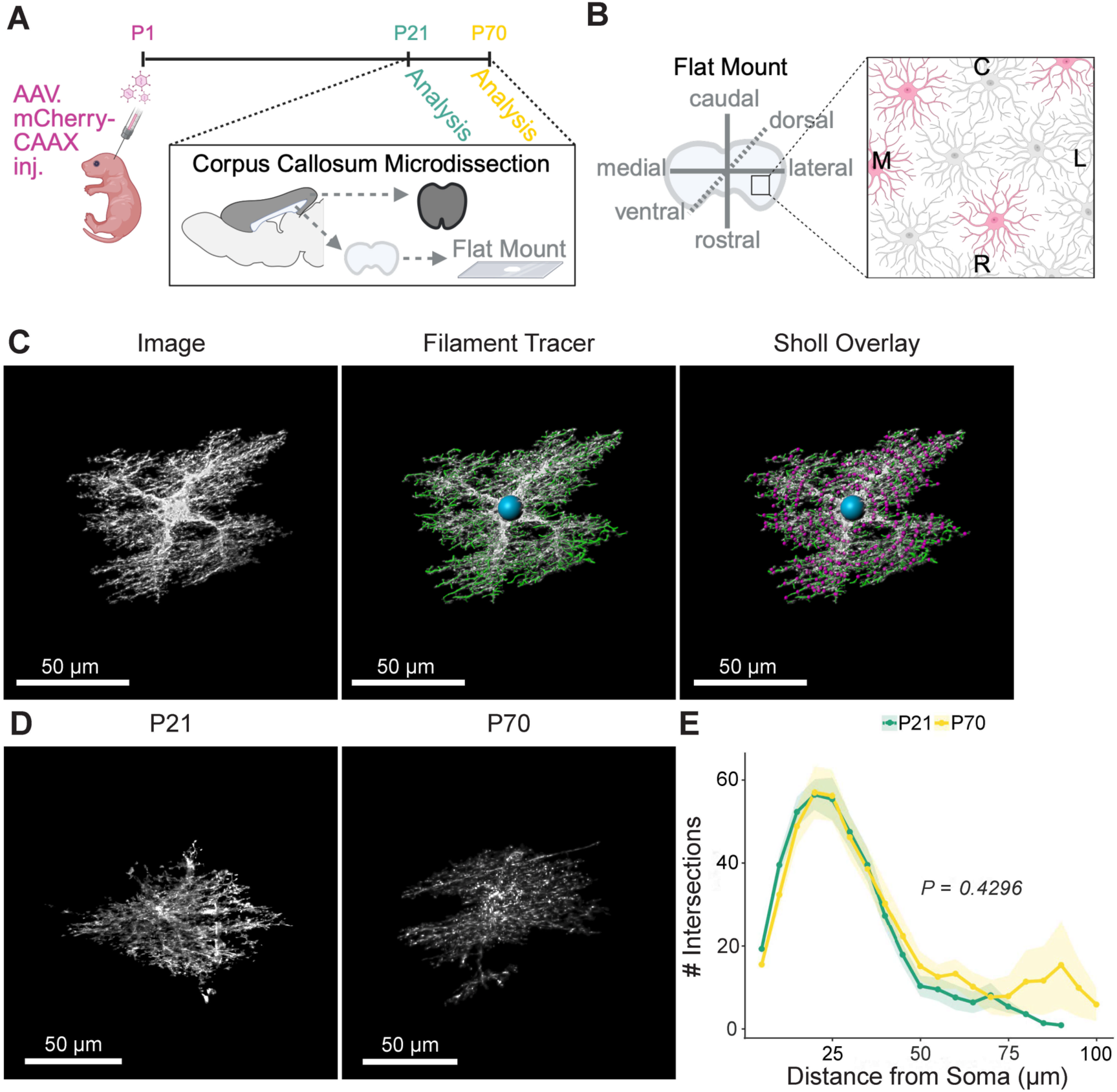
Branching complexity of WM astrocytes in unchanged between P21 and P70. **A)** Schematic of the AAV injection and the corpus callosum microdissection technique used to flat mount corpora callosi. Created with BioRender.com. **B)** Diagram showing that flat mounting the corpus callosum puts the rostral-caudal and medial-lateral axes in the x-y plane of the image. Created with BioRender.com. **C)** Representative example of the analysis pipeline which utilizes and IMARIS**-**based and AI-powered filament tracer to reconstruct WM astrocyte processes for *in vivo* Sholl analysis. **D)** Representative images of individual whole WM astrocytes expressing mCherry-CAAX in the flat mounted corpus callosum at P21 and P70. **E)** Sholl analysis of WM astrocytes branching complexity. Solid lines represent mean, and shaded areas represent ±SEM from 2-6 cells, from 4 animals at each timepoint and analyzed using a linear mixed model with Tukey HSD.

A comparison of branching complexity between P21 and P70 WM astrocytes revealed no significant difference between the two timepoints (Fig. 2D). The mean complexity of WM astrocytes at these two timepoints was similar and statistical analysis via mixed effects model revealed no significant difference between the two Sholl curves. In contrast to previously published studies that performed 3D Sholl analysis of GM astrocytes at similar time points and observed a Sholl curve peak near 300 (Coble et al., 2026; Tan et al., 2023), WM astrocytes are less complex, and their branches extend further from the soma. While we did not observe a significant increase in the mean complexity of WM astrocytes between P21 and P70, we did notice more intersections farther from the soma in P70 astrocytes, visible as a small peak around 90 µm (Fig. 2D). This peak reflects a sub-population of larger and more complex astrocytes that we observed at P70 and is consistent with the notably large variation in territory volume and surface area of P70 WM astrocytes in our morphology dataset (Fig. 1F, G).

### Astrocyte coverage of the corpus callosum increases into adulthood

We next asked whether the increase in WM astrocyte volume between P21 and P70 at the cellular level reflects an increase in astrocyte coverage throughout the corpus callosum. GM astrocytes carpet the cortex, and their coverage of the tissue increases as they morphologically develop (Stogsdill et al., 2017). Astrocytes in the corpus callosum are similarly distributed throughout the tissue but it’s unclear how this coverage changes during development. To confirm that astrocyte coverage increases with astrocyte size, we measured the coverage of WM astrocytes in the corpus callosum of Aldh1L1-eGFP mice at P8, P14, P21, and P60. Notably, Aldh1L1-eGFP labels astrocyte nuclei at all timepoints but is significantly dimmer in WM astrocyte processes at P60 than at the earlier timepoints (Supplemental Fig. 1A). This is consistent with transcriptomic profiling in adult mice that shows significantly lower Aldh1L1 expression in WM astrocytes compared to GM (Bocchi et al., 2025). The decrease in Aldh1L1-eGFP expression outside of the soma prevented our ability to accurately measure coverage using GFP signal at this timepoint. We therefore based our analysis of astrocyte coverage on the expression of GFAP which robustly labels the main branches of astrocytes in WM (Supplemental Fig. 1A). This approach introduces a limitation to this analysis in that it cannot account for additional coverage gained by the finer processes in the tissue yet ensures that comparisons can be made between timepoints as the same cellular marker is used. To evaluate astrocyte coverage, we acquired tiled confocal images to capture the continuous region of the corpus callosum below the motor and somatosensory cortex and performed analysis using a custom python-based FIJI macro. We first calculated the density of astrocytes within a manually drawn region of interest (ROI) containing the corpus callosum. We then counted the number of DAPI+ and Aldh1l1-GFP+ nuclei and divided this number by ROI area to calculate density. The process fraction was calculated by masking the GFAP channel in ImageJ and dividing the area of the GFAP+ processes by ROI area. The coverage index is the process fraction divided by the astrocyte density and reflects the area covered by each astrocyte.

As expected, the increase in astrocyte size seen between P21 and P60 coincided with an increase in astrocyte coverage (Supplemental Fig. 1B). Astrocyte density showed no significant differences between timepoints, though the average density at P60 was lower than at P21 (Supplemental Fig. 1C). This decrease in density could be an artifact of corpus callosum and WM astrocyte growth within the plane of the corpus callosum (the image z-plane), between these two time points. Because the size of our z-stack remains consistent at all time points, this would result in fewer nuclei captured within each image. The process fraction, or overall coverage, remained relatively consistent across the time points (Supplemental Fig. 1D). Similar to the astrocyte density, there was no significant difference across timepoints, but the average at P60 was lower than P21. As the astrocytes grow in the z-dimension, their major branches also spread out much like the nuclei, which may result in fewer branches captured in the z dimension at P60. Dividing the process fraction by a smaller number of astrocytes results in a larger area covered by each cell, suggesting that WM astrocytes expand between P21 and P70 to cover the corpus callosum.

### WM astrocytes tile at P14 and P21 in multiple dimensions, similar to GM astrocytes

As GM astrocytes grow in size and complexity to cover the cortex, they establish distinct, non-overlapping territories. This evolutionarily conserved phenomenon, known as tiling, is considered a hallmark feature of astrocytes across species (Bushong et al., 2002, 2004; Chen et al., 2020; Stork et al., 2014). Observational studies of WM astrocytes have suggested that they do not tile (Oberheim et al., 2009) but, to our knowledge, no published work has closely examined the tiling behavior of WM astrocytes in 3D. To address this knowledge gap and determine whether WM astrocytes share this hallmark feature with their GM counterparts, we conducted a thorough analysis of astrocyte tiling in WM astrocytes in multiple dimensions of the corpus callosum at both P14 and P21. Analysis of astrocyte tiling requires two adjacent astrocytes individually labeled with different fluorophores. To achieve this, we intracortically injected neonatal mice with two astrocyte-specific AAVs, one expressing GFP-CAAX and the other expressing mCherry-CAAX (Fig. 3A). We then collected images from coronal brain sections of neighboring astrocytes expressing different fluorophores in GM and WM at P14 and P21 and calculated the extent of territory overlap (Fig. 3B).

**Figure 3:**
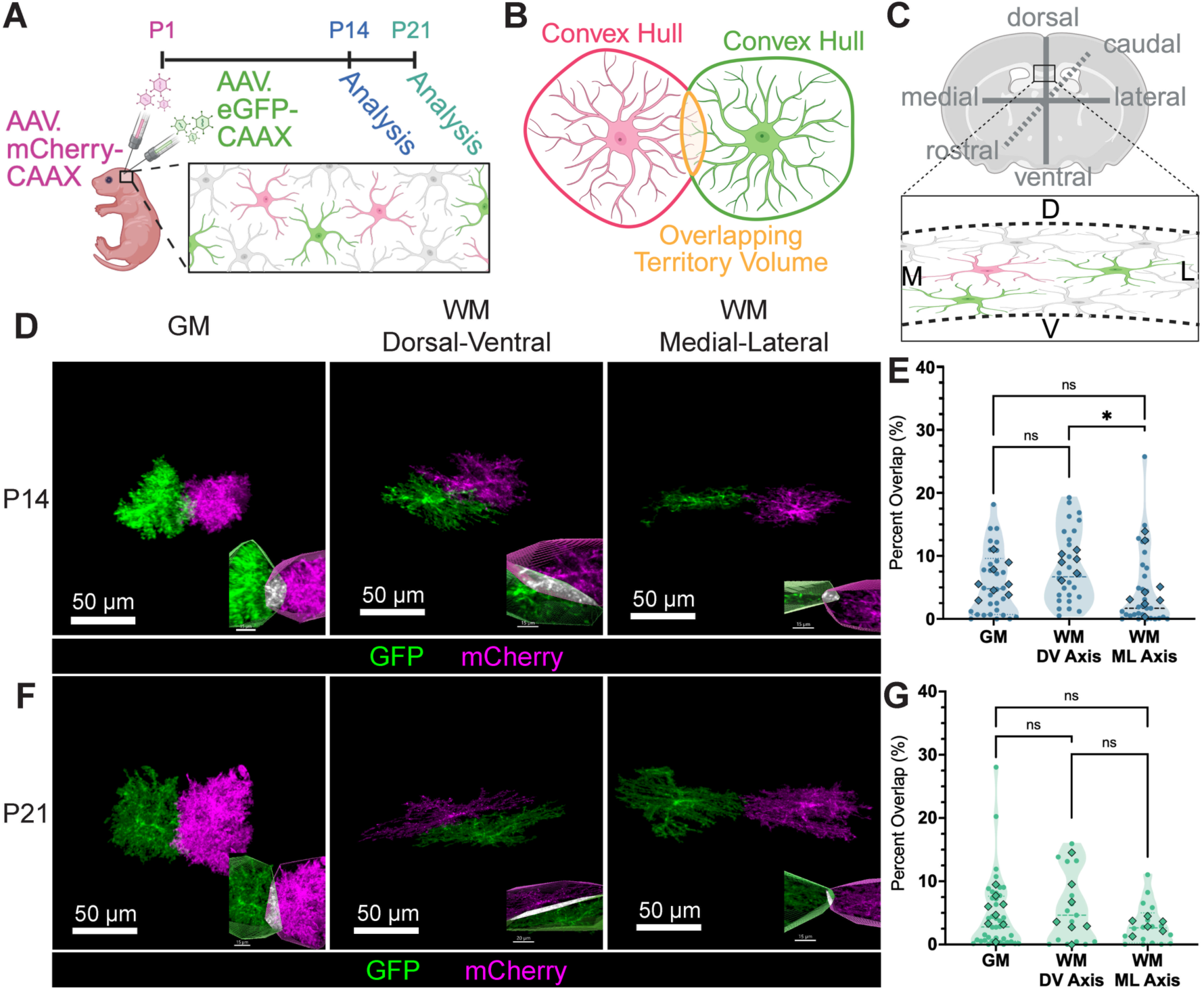
WM astrocytes tile at P14 and P21 in multiple dimensions, similar to GM astrocytes. **A)** Schematic of the AAV injection technique used to generate individual sparsely labeled astrocytes with two different fluorescent markers (GFP-CAAX and mCherry-CAAX) for tiling analysis. Created with BioRender.com. **B)** Schematic showing overlapping territory volume between two convex hulls generated in IMARIS. Created with BioRender.com. **C)** Schematic of the tiling dimensions represented in coronal section of the mouse corpus callosum. Created using BioRender.com. **D)** Representative confocal images of two whole astrocytes individually labeled with distinct fluorophores in GM and both dimensions of WM at P14. **E)** Violin plot comparing percent overlap between neighboring astrocytes in the cortex (GM), the dorsal-ventral axis of the corpus callosum (WM DV Axis), and the medial lateral axis of the corpus callosum (WM ML Axis) at P14. Dots represent individual cells and diamonds represent animal averages. 2-7 cells from 6-7 animals per condition, analyzed using a mixed effects model and one-way ANOVA with Sidak’s post-test for multiple comparisons. **F)** Representative confocal images of two whole astrocytes individually labeled with opposing fluorophores in GM and both dimensions of WM at P21. **G)** Violin plot comparing percent overlap between neighboring astrocytes in the cortex, the dorsal-ventral axis of the corpus callosum, and the medial lateral axis of the corpus callosum at P21. Dots represent individual cells and diamonds represent animal averages. 1-7 cells from 6-8 animals per condition, analyzed using a mixed effects model and one-way ANOVA with Sidak’s post-test. ns = p<0.05, * = p≤0.05, ** = p≤0.01, ***= p≤0.001, **** = p≤0.0001.

Astrocytes in GM occupy territories that are largely spherical or ellipsoid, conferring a relatively equal capacity for tiling in different anatomical axes. WM astrocytes in the corpus callosum are flattened in the dorsal-ventral axis and occupy circular or elliptical territories in the medial-lateral and rostral-caudal axes. As a result, neighboring astrocytes which are stacked in the dorsal-ventral axis of the corpus callosum have a larger cellular interface than neighboring astrocytes in the medial-lateral or rostral-caudal axis (Fig. 3C, Fig. 2B). To reflect the asymmetry of these cells, we assessed WM astrocyte tiling in both dorsal-ventral and medial-lateral axes. Of note, we did not assess tiling in the rostral-caudal axis due to technical limitations, as this dimension is restricted by the thickness of coronal sections. To measure tiling, we calculated the volume of overlapping territory and dividing this by the territory volume of the larger astrocyte to obtain the percent overlap (Supplementary Video 1). We found that the percent overlap between WM astrocytes in either dimension is not significantly different from GM astrocytes at P14 or P21 (Fig. 3E, G). At P14, we found that WM astrocytes have a higher percent overlap in the dorsal-ventral axis than in the medial-lateral axis which likely reflects the large cellular interface that neighboring astrocytes share in this axis. Collectively, these results demonstrate that WM astrocytes, like GM astrocytes, establish non-overlapping territories and tile the corpus callosum at P14 and P21.

### Most, but not all, mature WM astrocytes have endfeet

Contact with vasculature is a hallmark feature of GM astrocytes. In mouse cortical GM, all astrocytes contact the vasculature with at least one endfoot process, and the extent of vascular contact appears to scale with blood vessel density (Hösli et al., 2022). WM is less vascularized than GM (Cavaglia et al., 2001) and though WM astrocytes are known to have endfeet (Gleichman et al., 2025), whether contact with vasculature is a hallmark feature of WM astrocytes is unknown. To qualitatively assess individual WM astrocytes for endfeet, we stained tissue with sparsely labeled mCherry-CAAX astrocytes (Fig1. A) for aquaporin 4 (AQP4), a water channel that is heavily enriched at astrocyte endfeet. We imaged whole astrocytes in thick tissue sections via confocal microscopy and identified endfeet in IMARIS 3D viewer using a combination of their distinct tubular morphology, AQP4 expression, and co-localization of AQP4 and mCherry-CAAX (Fig. 4A, B). Astrocytes with at least one endfoot were categorized as “present” while astrocytes with no visible endfoot were categorized as “absent.” We performed this analysis at the same four timepoints as the morphology analysis shown in Fig1: P8, P14, P21 and P70. Our analysis revealed that while most WM astrocytes have endfeet, some do not. We found that the percentage of WM astrocytes with endfeet increases over development (Fig. 4C). This increase over development is statistically significant (Chi-Squared p=0.00174) and the standard residuals indicate that there are fewer than expected astrocytes with endfeet at P8 (r =-3.58) and more than expected at P21(r =2.49) which indicates that endfeet largely develop between P8 and P21 (Fig. 4C). By P21, 94.4% of the astrocytes we analyzed had endfeet. At P70, 91.3% of astrocytes had endfeet suggesting that astrocytes in the corpus callosum do not continue developing endfeet into adulthood. These results indicate that while contact with vasculature is a common feature of WM astrocytes, there is a subpopulation of WM astrocytes with no apparent vascular contact.

**Figure 4:**
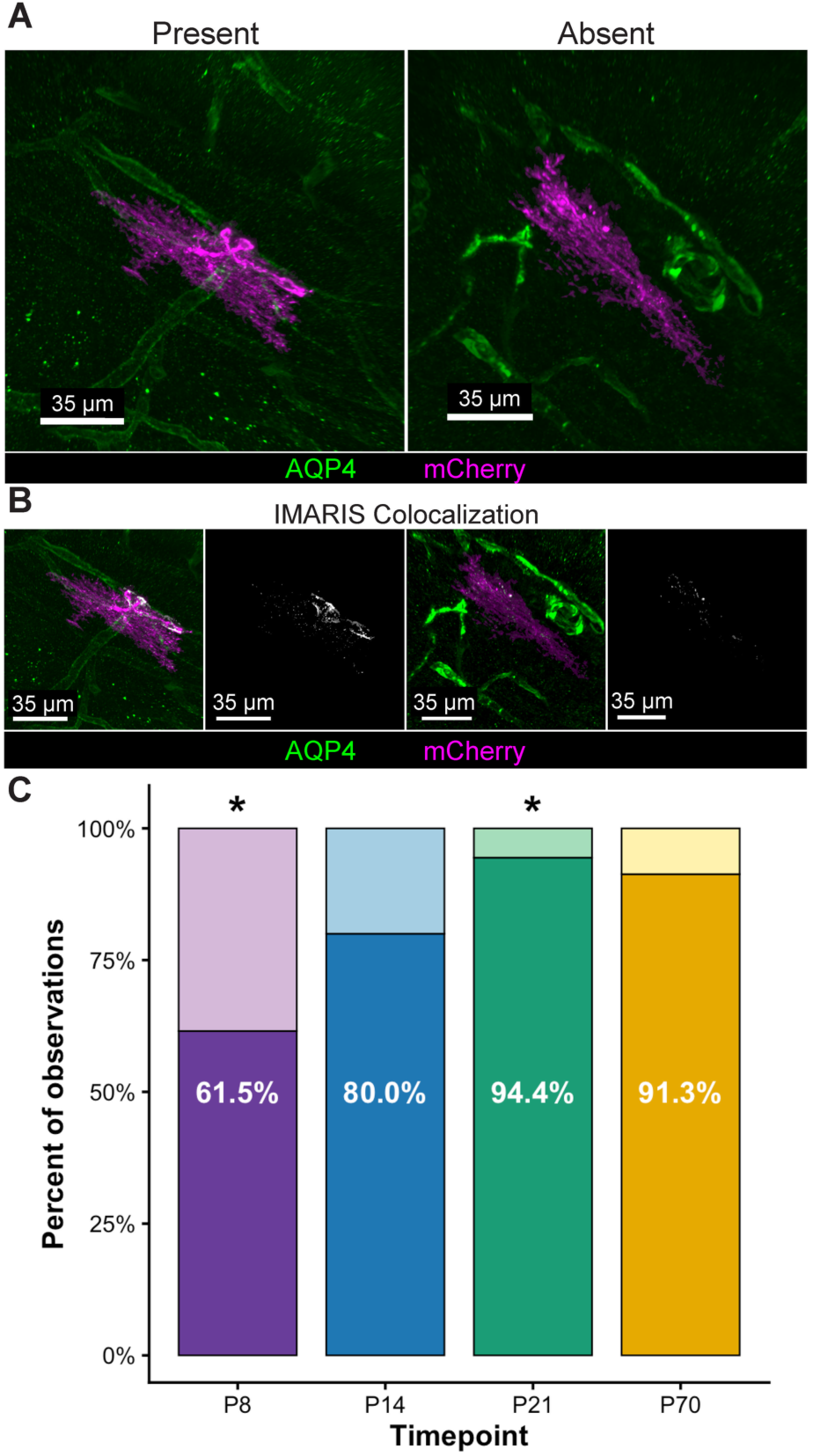
Most, but not all, mature WM astrocytes have endfeet. **A)** Representative confocal images of whole WM astrocytes with (present) and without (absent) endfeet at P21. **B)** IMARIS analysis of the colocalization between the astrocyte (mCherry-CAAX, magenta) and AQP4 (green) to visualize endfeet in white. **C)** Stacked bar graph for the frequency of present and absent endfoot observations at each timepoint in the corpus callosum. 41, 30, 36, and 23 cells were analyzed at P8, P14, P21 and P70 respectively. Cells were sampled from 6-7 animals per timepoint. Chi-squared test of independence with standard residuals. ns = p<0.05, * = p≤0.05, ** = p≤0.01, ***= p≤0.001, **** = p≤0.0001.

### WM astrocytes closely associate with axons at early timepoints

Our findings thus far indicate that astrocytes in WM share some characteristic features with their GM counterparts including endfeet and tiling behavior but exhibit notable differences in their morphology and developmental trajectories. Close association with neuronal synapses is another hallmark feature of GM astrocytes and interaction with synapses drives the morphological maturation of GM astrocytes (Baldwin et al., 2024; Cheng et al., 2023; Stogsdill et al., 2017). The WM microenvironment is axon-rich and largely devoid of neuronal synapses, though axons do form synaptic connections with oligodendrocyte precursor cells (Bergles et al., 2000). To quantitatively compare the synaptic environment of WM astrocytes to GM, we compared the density of excitatory synapses and pre and post synaptic markers in layer 5 of the somatosensory cortex and the corpus callosum at P14, P21 and P60 in Aldh1L1-eGFP mice (Supplemental Fig. 2A). We identified synapses by colocalization of the excitatory presynaptic marker VGlut1 and the excitatory postsynaptic marker PSD95. Using Synbot (Savage et al., 2024), we calculated the density of co-localized puncta (Supplemental Fig. 2B), VGlut1 puncta (Supplemental Fig. 2C), and PSD95 puncta (Supplemental Fig. 2D). As expected, the density of synapses in the WM was substantially lower than GM at all three timepoints (Supplemental Fig. 2B) as was the density of VGlut1 and PSD95 puncta (Supplemental Fig. 2C, D). These results indicate that the corpus callosum is a synapse-poor environment, suggesting that the morphological maturation of WM astrocytes is likely driven by synapse-independent mechanisms.

Axons are the prominent structural component of WM during development. Pioneer callosal axons begin traversing the corpus callosum at E15.5 and other axons follow around E17.5 (De León Reyes et al., 2020; Gavrish et al., 2024). The precise timing of axon development in the corpus callosum has proven difficult to establish, but evidence suggests that the number of callosal axons increases until P15 (Mizuno et al., 2007) and declines due to pruning by P30 (De León Reyes et al., 2019). This suggests that axons are still transversing the corpus callosum during the early window of WM astrocyte morphogenesis between P8-P14. To understand the spatial arrangement of axons and WM astrocytes during this time frame, we stained tissue with sparse mCherry-CAAX astrocyte labeling at P7 and P12 with the axonal marker TUJ1. At P7, axons appear to be more disorganized with astrocyte branches extending more radially. By P12, axons are more structured with astrocyte processes organized along and within axon bundles (Fig 5A). Collectively, these images reveal that astrocytes closely associate with axons during early stages of morphological maturation, layering themselves within developing bundles of axons and orienting their processes in parallel to axons.

**Figure 5:**
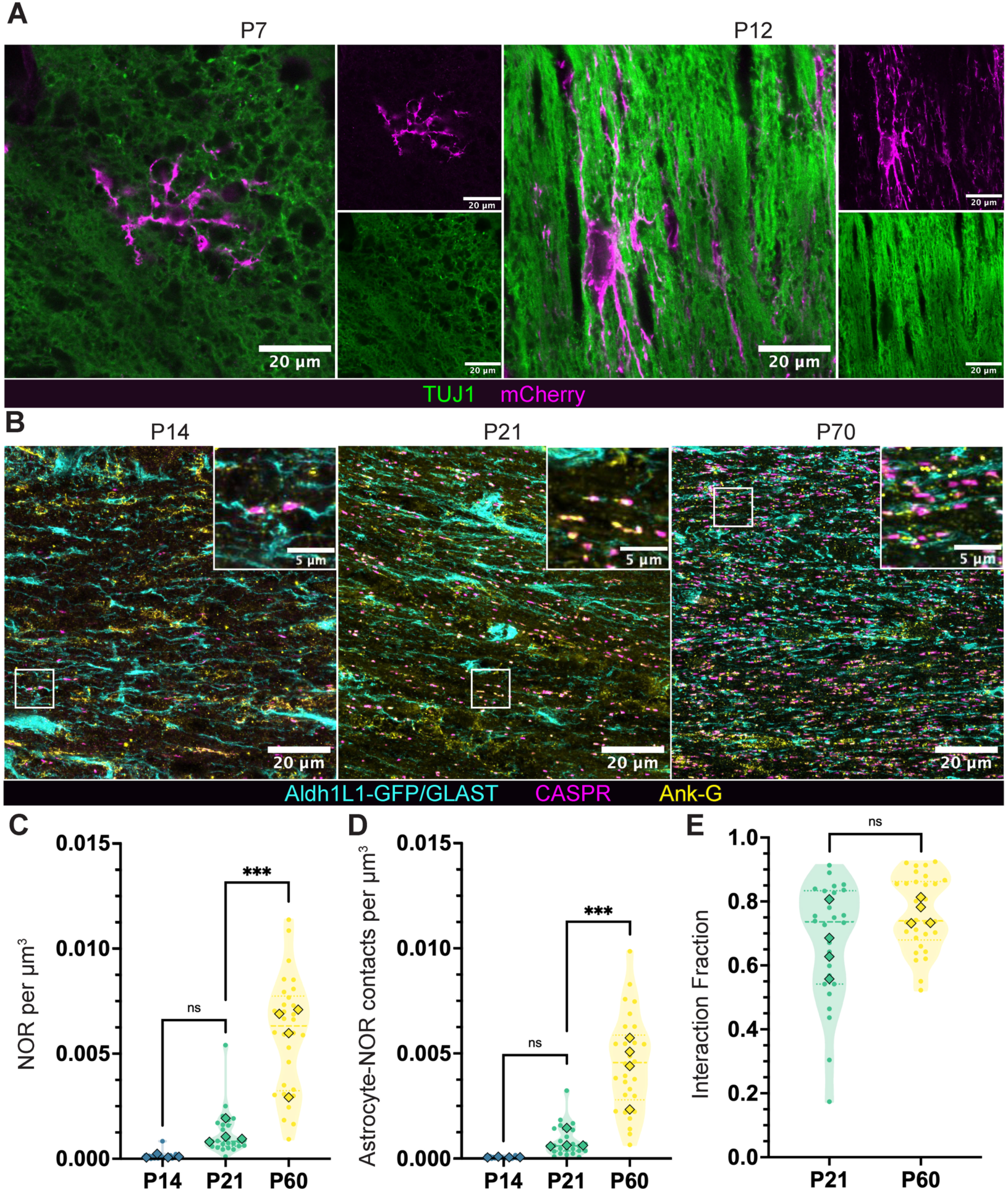
Node of Ranvier formation in the corpus callosum temporally coincides with morphological maturation in WM astrocytes. **A)** Representative confocal images of individually labeled astrocytes (mCherry-CAAX, magenta) and axons (TUJ1, green) at P7 and P12. **B)** Representative confocal images of astrocyte processes, as labeled by Aldh1L1 (cyan, P14 and P21) or Aldh1L1 and GLAST (cyan, P60), and Nodes of Ranvier labeled by the combination of CASPR (magenta) and Ank-G (yellow) in the corpus callosum. Insets, highlighted on the full images with white boxes, show examples of astrocyte-node interactions. **C-E)** Violin plots of **C)** Node of Ranvier density (NOR per µm^3^) **D)** Astrocyte-Node of Ranvier interactions per µm^3^ (Astrocyte-NOR contacts per µm^3^) at P14, P21, and P60 and **E)** interaction fraction, percentage of NORs with astrocyte contact at P21 and P60. For all plots, dots represent individual images and diamonds represent animal averages. 3-8 images per animal from 4 animals were analyzed at each timepoint. Nested one way ANOVA with Sidak’s post-test. ns = p<0.05, * = p≤0.05, ** = p≤0.01, ***= p≤0.001, **** = p≤0.0001.

### Node of Ranvier formation in the corpus callosum temporally coincides with morphological maturation in WM astrocytes

This within-bundle orientation of astrocytes in the corpus callosum positions astrocytes to communicate along the length of axons. As myelination progresses during development, Nodes of Ranvier (NOR) form at gaps in the myelin sheath to regulate signal propagation down the axon. WM astrocytes are known to contact NOR to regulate axon conductance and ion homeostasis. (Dutta et al., 2018; Hamilton et al., 2008; Kalsi et al., 2004; Lezmy et al., 2021). The developmental timeline of NOR establishment in the corpus callosum has not been thoroughly characterized, but likely coincides with the onset of myelination, which begins around P8 in this region and continues through adulthood (Ozarkar et al., 2024). To determine whether NOR establishment in the corpus callosum temporally coincides with the morphological maturation of WM astrocytes, we measured the density of NOR and NOR-astrocyte interactions in the corpus callosum at P14, P21 and P60. To do so, we performed immunolabeling in coronal sections of the medial corpus callosum under the somatosensory and motor cortexes in Aldh1L1-eGFP mice to detect CASPR, a paranodal transmembrane protein which links the axonal membrane to the myelin sheath, and Ankyrin-G (AnkG), a scaffolding protein that anchors ion channels at the center of the node. Dual labeling with CASPR and AnkG produces a robust dash-dot-dash pattern around NOR (Fig. 5B). Because Aldh1L1-eGFP is weakly expressed in astrocytes branches at P60 (Supplemental Fig. 1A), we co-stained the P60 tissue for GLAST which robustly labels the finer process of WM astrocytes in adulthood (Fig. 5B). Because GLAST expression has also been reported in OPCs, we stained P60 tissue for GFP, GLAST, and PDGRFa which labels OPCs. We found minimal colocalization between GLAST and PDGRFa and robust colocalization between major astrocyte branches labeled with GFP and GLAST (Supplemental Fig. 3A). Without GLAST, interactions between NOR and fine astrocyte processes in adulthood are not apparent (Supplemental Fig. 3B, C).

To calculate NOR density, we developed a machine learning powered analysis pipeline in IMARIS 11 trained to recognize the dash-dot-dash pattern of the CASPR and AnkG in 3D regions of interest (ROIs) (Supplemental Fig. 4). To identify astrocyte-NOR interactions, we constructed surfaces of astrocytes branches within the ROI and filtered the AnkG spots within nodes for proximity to the astrocyte surface. Though myelination is reported to begin around P8 in the corpus callosum, electron microscopy analysis reveals that approximately 10% of axons are myelinated by P30 (Ozarkar et al., 2024). Consistent with this, NOR density remained low at P14 and P21 (Fig. 5C), a timepoint by which WM astrocytes have already undergone a significant increase in morphological complexity (Fig. 1E). Between P21 and P70, a time frame during which WM astrocytes grow significantly in volume but not complexity, we observed a robust increase in NOR density (Fig. 5C), coinciding with the progression of myelination (Sturrock, 1980). Similarly, the density of astrocyte-NOR interactions increased significantly between P21 and P70 (Fig. 5D). Of note, the proportion of NOR which interact with astrocytes remained consistent between P21 and P70, at approximately 75% for both timepoints (Fig. 5E). This interaction fraction, or the percentage of NOR with an interacting astrocyte, was not calculated at P14 because the exceedingly low density of NOR made it mathematically impractical. These results indicate that between P21 and P70 there is a significant increase in the density of NORs and astrocyte-NOR contacts in the corpus callosum, concurrently with significant growth of WM astrocytes.

## Discussion

In this study, we conducted a thorough analysis of the morphological features of WM astrocytes in the corpus callosum across postnatal mouse development. Our gross morphological analysis comparing cortical GM astrocytes to callosal WM astrocytes highlights their unique shapes. As expected, WM astrocytes are less spherical and less complex than their GM counterparts at all timepoints. Interestingly, mature astrocytes have similar surface area in both tissues suggesting that these two types of astrocytes may have a similar maximum supportable membrane. Our analysis also provided previously unknown insight into the developmental trajectories of WM astrocytes. Astrocytes in the corpus callosum increase in branching complexity early in development from P8 to P14 and subsequently increase significantly in volume from P21 to P70. In contrast, cortical astrocytes increase in volume from P8 to P14 while progressively increasing branching complexity from P8 to P21. Our data indicates that WM astrocytes take longer to reach morphological maturity than those in GM and highlights that astrocytes in these different environments exhibit distinct developmental trajectories.

While our morphological analysis highlights the differences between GM and WM astrocyte structure, we also find common features between GM and WM astrocytes that are conserved not only throughout the brain, but across evolution (Chen et al., 2020). Astrocyte tiling is observed in Drosophlia (Stork et al., 2014), zebrafish (Chen et al., 2020), rodent (Bushong et al., 2002, 2004), and human (Oberheim et al., 2009), though human astrocytes show a greater degree of territory overlap. The physiological function of tiling is unclear, but this feature is often disrupted in disease (Li et al., 2019; Oberheim et al., 2009) and is likely important for astrocyte network organization (Baldwin et al., 2021). A previous study suggested that WM astrocytes do not tile, though this observation was based on a 2D image of GFAP immunolabeling in human tissue where cellular domains are difficult to distinguish and processes appear to overlap (Oberheim et al 2009). In our analysis, we directly measured tiling in 3D images of neighboring WM astrocytes in the corpus callosum that expressed unique fluorophores. We found that astrocyte tiling is conserved in this WM tract at P14 and P21 and is present along multiple anatomical axes. To our knowledge, ours is the first study to directly measure tiling in callosal WM astrocytes and indicates that tiling is a shared feature of GM and WM astrocytes. Notably, we did not examine whether this tiling persists into adulthood. Our morphology data shows that WM astrocytes increase significantly in size between P21 and P70 and this increase in size may require astrocytes to encroach on each other’s territory. Alternatively, corpus callosum growth during this time frame and into adulthood might allow astrocytes to expand their territories while maintaining tiling.

Vascular contact via specialized endfeet processes is another conserved feature of astrocytes in species with a vascular network (Gall et al., 2025; Hösli et al., 2022). In the mouse brain, all cortical astrocytes contact vasculature with at least one endfoot. In deeper cortical layers, where blood vessel density is higher, astrocytes contact more vessels. In the hippocampus, which is less vascularized than the cortex, some astrocytes do not contact vasculature (Hösli et al., 2022). Here, we found that most mature astrocytes in the corpus callosum have endfeet, but a subset do not. WM is known to have lower blood vessel density than GM (Cavaglia et al., 2001), which may mean that some astrocytes occupy territories that do not include a blood vessel and, as a result, do not develop endfeet.

The morphological differences between GM and WM astrocytes, from distinct geometry to variations in vascular contact, likely reflect their vastly different microenvironments. GM astrocytes reside in a dense network of synapses, neuronal cells bodies, unmyelinated axons, dendrites, and glial cell processes. In contrast, WM is largely devoid of neuronal cell bodies and synapses and is instead rich in axons and OPCs in early postnatal development, with a progressive increase in mature oligodendrocytes and myelin towards adulthood. Direct interaction with neuronal synapses is a major driver of GM astrocyte morphogenesis, yet WM has very few synapses. In an environment devoid of synapses, how do astrocytes develop? Given that GM astrocyte morphogenesis depends heavily on cues in the immediate microenvironment, the cellular composition of the developing corpus callosum, in concert with the developmental trajectory of WM astrocytes that we describe in this paper, may offer some clues.

We show that WM astrocytes develop in a biphasic fashion, with the early phase consisting of increases in branching complexity (P8-P14) and the later phase consisting of increases in territory volume (P21 – P70). This developmental trajectory mirrors two major phases of corpus callosum development: callosal axon development and myelination. Pioneer callosal axons begin transversing the corpus callosum at embryonic day (E) 15.5 and other axons follow around E17.5 (De León Reyes et al., 2020; Gavrish et al., 2024). Current evidence suggests that the number of callosal axons peaks around P15 and are subsequently pruned until around P30 (De León Reyes et al., 2019; Mizuno et al., 2007). Myelination in the corpus callosum begins around P10. By P30, approximately ten percent of corpus callosum axons are myelinated and myelination increases substantially into adulthood (Ozarkar et al., 2024). In sum, corpus callosum development is biphasic much like the maturation of its resident astrocytes, with the early phase (E15.5-P15) dominated by axon growth and the later phase (juvenile to adulthood) by myelination. During the early phase, we found a close association between WM astrocytes and axons, while the later phase showed a stark increase in astrocyte-NOR association. Based on these observations, we propose a working model of WM astrocyte development, wherein the early phase of maturation is guided primarily by interactions with developing axons and the later phase of development is driven by interactions with differentiating oligodendrocytes and myelin (Supplemental Fig. 5). In addition to environmental cues, other factors may contribute to WM astrocyte development and could expand upon this model in the future. For example, though strong evidence establishes tissue microenvironment as a key contributor to structural and molecular heterogeneity in astrocytes (Endo et al., 2022), some differences between GM and WM astrocytes could arise from differences in developmental origin, as clonal analysis shows regional variation in the source of progenitors (Ojalvo-Sanz et al., 2024; J. Zhou et al., 2025). Moreover, the directional arrangement of axonal tracts imposes structural constraints and differences in other physical properties such as tensile strength which could impact cell structure.

Our findings also inspire questions about heterogeneity within WM astrocytes. In particular, we found that mature callosal astrocytes display large variability in territory volume suggesting that there is morphological heterogeneity among astrocytes in the corpus callosum. While our study focused only on the corpus callosum, heterogeneity between WM astrocytes associated with different regions of the optic tract has been established (Holden & Calkins, 2025). The mechanisms underlying astrocyte heterogeneity are an active area of investigation and there is evidence that heterogeneity may be developmentally encoded both temporally and spatially through a cell’s lineage (Ojalvo-Sanz et al., 2026) and arise from a cell’s local environment (Endo et al., 2022). The spatial arrangement of WM tracts in the rodent brain highlights the need for high-resolution, high-throughput, and volumetric imaging modalities capable of comparing individual astrocyte morphology across many microenvironments and brain regions. This kind of imaging would allow us to understand both within and between tissue variation among astrocytes both in WM and GM.

Whether WM astrocyte morphology differs in species with higher WM content, such as humans and non-human primates, is unknown. We know that human WM astrocytes share a similar molecular signature to their murine counterparts (Bocchi et al., 2025) and that human WM astrocytes resemble those in the mouse in shape but are larger in diameter (Oberheim et al., 2009). However, no studies have yet assessed WM human astrocyte morphology in detail, nor have they explored heterogeneity in human astrocytes. Studies of astrocyte morphology in human tissues primarily use GFAP to label astrocytes in thin tissue sections (Man et al., 2024; Oberheim et al., 2009). This strategy does not capture the full branching complexity or the full cell volume, making comparisons of WM astrocyte complexity between species challenging. A recent study that examined human cortical GM and WM tissue via 3D electron microscopy does offer some insight into human WM astrocyte morphology (Shapson-Coe et al., 2024) and may provide a starting point for understanding similarities and differences between rodent and human WM astrocytes. The extent of WM astrocyte heterogeneity in humans and whether unique WM astrocytes subtypes exist in human WM as they do in human GM is also unknown. A significant challenge to studying human WM astrocytes is the lack of faithful *in vitro* models, like 3D culture systems and organoids, capable of mimicking the structure and composition of WM tracts in any species. Generating novel tools to overcome these limitations would greatly improve our ability to answer questions about WM astrocyte development and function in humans and in turn, would expand our understanding of the role of WM astrocytes in disease.

WM astrocyte dysfunction plays a causative role in leukodystrophies such as Alexander Disease and Vanishing White Matter Disease (Dooves et al., 2016; Leferink et al., 2019; Sosunov et al., 2018; L. Zhou et al., 2019). Disorders such as multiple sclerosis disproportionally affect WM, and most other brain disorders from Alzheimer’s Disease to Autism Spectrum Disorder feature WM pathology (Davis et al., 2003; Fitzgerald et al., 2019; Frischer et al., 2015; Nasrabady et al., 2018; Ozarkar et al., 2024). Thus, a detailed understanding of WM astrocyte development and function is a critical step to understanding and treating WM dysfunction. Our study lays the foundation for further study of WM astrocytes in models of neurological disorders.

## Author Contributions

K.H.M.H.: conceptualization, methodology, investigation, formal analysis, visualization, writing – original draft. M.F.: investigation, methodology, formal analysis, visualization, writing – review and editing. K.T.B: conceptualization, methodology, investigation, formal analysis, supervision, funding acquisition, visualization, writing – original draft.

## Acknowledgments

We thank Baldwin lab members Amy Stanek, Hayli Spence-Osorio, and Elliot Evans for their assistance in generating and processing samples used in the study.

## Methods

### Animals

All mice were used in accordance with the Institutional Animal Care and Use Committee (IACUC) and the UNC Department of Comparative Medicine (IACUC Protocol Number 24-005). Mice were housed in standard conditions with 12-hour day/night cycles. Timed-pregnant CD1 females were obtained from Charles River (RRFD:IMSR_CRL:022) and Aldh1l1-GFP transgenic mice were obtained from MMRRC (RRID:MMRRC_011015-UCD).

Mice were collected for experiments between postnatal day 8 (P8) and P70 as specified in the text and figure legends. For all experiments, mice of both sexes were included in analysis, and we did not observe any influence or association of sex on the experimental outcomes. Criteria for inclusion, exclusion, and randomization are listed for each experiment in specific methods subsections.

### AAV Production and Administration

The pZac2.1-gfaACB1D plasmids containing gfaABC1D-GFP-CAAX (Baldwin et al., 2021) or gfaABC1D-mCherry-CAAX (Stogsdill et al., 2017) were packaged into AAV2/5/PHP.eB capsids by the UNC BRAIN Initiative Viral Vector Core as previously described. Briefly, purified AAVs were exchanged into storage buffer containing 1 x phosphate-buffered saline (PBS), 5% D-Sorbitol, and 350 mM NaCl. Virus titers (GC/ml) were determined by qPCR targeting the AAV inverted terminal repeats. All viruses were adjusted to the same titer (3.4×10^13^ GC/mL) and 1 μL of virus was injected unilaterally into the cortex of hypothermia-anesthetized postnatal day 1 (P1) CD1 mice using a Hamilton syringe with a custom removeable needle. For morphology studies, including corpus callosum flat mount, only the mCherry-CAAX virus was injected unilaterally. For tiling studies, the mCherry-CAAX and GFP-CAAX viruses were injected unilaterally on opposite sides.

### Immunohistochemistry

#### Coronal Section Sample Preparation

Brains were collected, frozen, sectioned, and stained as described previously (Eaker & Baldwin, 2022) Breifly, mice were anesthetized with 0.8 mg/kg tribromoethanol (avertin) and perfused with TBS/Heparin, followed by 4% PFA in TBS. Brains were post-fixed overnight in 4% PFA in TBS and cryoprotected in 30% sucrose in TBS for at least 2 days. Brains were frozen in embedding molds using a medium containing two parts 30% sucrose in TBS and one-part O.C.T. (Tissue-Tek) and stored at -80 °C. Frozen brains were sectioned coronally to 40 µm or 100 µm thickness using a CryoStar NX50 Cryostat (Thermo Fisher Scientific) and stored in a 1:1 mixture of glycerol and 1xTBS at -25 °C until use. For immunostaining, all washes and incubations were performed on a platform shaker at 100 rpm. Sections were washed in 3 x 10 minutes in TBS containing 0.2% Triton (TBST) and blocked in blocking solution (10% goat serum in TBST) for 1 hour at room temperature. Primary antibodies were diluted in blocking solution and sections were incubated for 2 nights (40 µm) or 3 nights (100 µm) at 4 °C while (see specific sections for primary antibody information and concentration). Following primary antibody incubation, sections were washed in 3 x 10 minutes in TBST, incubated in secondary antibody solution (1:200 in blocking solution) for 2 hours (40 µm) or 3 hours (100 µm) at room temperature and washed 3 x 10 minutes in TBST. For some experiments, DAPI was added to the secondary antibody solution for the final 10 minutes of incubation at a 1:50,000 concentration. Sections were transferred to a 2:1 mixture of TBS:H2O and then mounted onto glass slides with homemade mounting medium (20 mM Tris pH 8.0, 90% Glycerol, 0.5% N-propyl gallate), sealed with nail polish, dried at room temperature, and stored at 4 °C until imaging. For primary antibodies produced in mouse, isotype specific secondary antibodies were used (e.g., goat anti-mouse IgG1) to prevent excessive background staining.

#### Corpus Callosum Flat Mount Sample Preparation

CD1 mice with unilateral AAV-mCherry-CAAX were perfused at P21 or P70 as described above, post-fixed overnight in 4% PFA, and stored in TBS + 0.02% sodium azide at 4 °C before microdissection. Microdissection was performed in cold TBS using a dissection microscope (Olympus SZX-ZB7), a clean razor blade, and Roboz Surgical Super Fine #5SF tweezers. First, the corpus callosum was exposed by removing the cerebellum and occipital lobes with the razor blade. The razor blade was then used to cut the brain along the midline producing two hemispheres. The midbrain was carefully removed using two pairs of tweezers to precisely cut away tissue. As the corpus callosum became visible, the ventral most edge of the cortex was cut diagonally as this section of the corpus callosum is too thin to remove intact. The tweezers were then run along the border between the cortex and corpus callosum to separate the two tissues. Once separation was complete, the corpus callosum was peeled away and divided into 2 pieces. This process was then repeated for the other hemisphere, and samples were stored in 1:1 mixture of glycerol and TBS at -25 °C until use.

Flat mount samples were stained using the above protocol with increased time in blocking (4 hours), primary (4 nights) and secondary (4-5 hours) to allow antibodies time to penetrate the thicker tissue. DAPI was added to the secondary antibody solution for the final 10 minutes of incubation at a 1:50,000 concentration. Glass slides were prepared with label tape spacers to prevent crushing the corpus callosum. Corpora callosa were mounted onto prepared glass slides with homemade mounting medium (20 mM Tris pH 8.0, 90% Glycerol, 0.5% N-propyl gallate) and sealed with nail polish.

### 3D Astrocyte Morphology Analysis

Gross morphology of individual astrocytes was assessed in 100 μm-thick coronal sections containing motor and somatosensory cortex and the corpus callosum. Tissue from CD1 mice previously intracortical injected with mCherry-CAAX AAV was collected at postnatal day 8 (P8), P14, P21, and P70. Immunohistochemistry (IHC) was performed using rat anti-RFP primary antibody (1:1000, Thermo Fisher, M11217) to detect mCherry and guinea pig anti-AQP4 (1:500, Alomone Labs, AQP-014-GP), followed by goat anti-rat Alexa Fluor 594 secondary antibody (1:200, Thermo Scientific, A-11007) and goat anti-guinea pig Alexa Fluor 488 secondary antibody (1:200, Thermo Scientific, A-11073). High magnification confocal z-stack images (1024 x 1024) containing whole astrocytes were acquired on an Olympus FV3000 microscope with a 30x objective and 2x optical zoom with a 0.5 µm step size. Z-stack size was adjusted to ensure the full astrocyte was captured. Gray matter (GM) astrocytes images were collected from layers 2/3 through 5 of motor and somatosensory cortexes and white matter (WM) astrocyte images were collected from the medial corpus callosum. We defined this region as the continuous corpus callosum underneath the primary and secondary motor cortexes and the somatosensory cortex medial to the barrel cortex. Images of this region can be found in Supplemental Figure 1. Astrocytes that were not fully contained within the section or were not located in the specified regions were not imaged. Imaris 10.2 (Bitplane) software was used as to measure surface area, territory volume, and sphericity. Minimal post-processing (Median Filter 3×3×3) was performed in Imaris to aid whole-cell surface reconstruction with the surface creation tool. Spots close to surfaces were then generated and used to build a convex hull using a custom MATLAB Convex Hull Xtension around the whole astrocyte territory as described previously (Eaker & Baldwin, 2022). Astrocytes were excluded from the final analysis if exceptionally dim fluorescent signal or image poor signal to noise ratio prevented accurate surface generation or if the boundaries of the astrocyte territory could not be distinguished. CSV Files containing detailed surface and convex hull measurements were exported from Imaris using the Vantage function. The territory volume was obtained from the volume of the convex hull and represents the volume of the anatomical region occupied by the astrocytes. Relative Complexity measures how much of the cell’s territory volume is filled by the astrocyte and its processes and is calculated by dividing volume of the surface generated from the mCherry-CAAX labeling by the territory volume. Surface area measures the area of the covering the surface volume and estimates the area of the cell’s membrane. Sphericity is calculated from the convex hull and describes how close the cell’s shape is to a sphere. Mathematically, a sphericity of 0 is a two-dimensional dot while 1 is a perfect sphere such that all 3D shapes have a sphericity of at least 0.5.

Statistical analysis was performed in GraphPad Prism 11 (Version 11.0.2). Individual cell measurements for territory volume, surface area, relative complexity, and sphericity were obtained from 3-9 cells per animal from 4-7 animals for both GM and WM astrocytes at 4 developmental timepoints (P8, P14, P21, and P70). Animals with fewer than three analyzable cells were excluded to ensure reliable estimation of animal-level effects. Normality was assessed using the Shapiro–Wilk test and visual inspection of QQ plots. Differences among experimental groups were evaluated using a one-way nested ANOVA with Sidak’s post-test, with cells nested within animals and animals nested within timepoint and tissue type conditions. Graphs display individual cell measurements and animal-level means. The number of animals and cells/animal analyzed is indicated in the corresponding figure legend for this experiment. Statistical significance was defined as *P* < 0.05.

### In vivo Sholl Analysis

*In vivo* Sholl analysis was performed in flat mount corpus callosum. IHC was performed as described above using rat anti-RFP primary antibody (1:1000, Thermo Fisher, M11217) and goat anti-rat Alexa Fluor 594 secondary antibody (1:200, Thermo Scientific, A-11007) and DAPI. High magnification confocal z-stack images (1024 x 1024) containing whole astrocytes were acquired on an Olympus FV3000 microscope with a 30x objective and 2x optical zoom with a 0.5 µm step size. Z-stack size was adjusted to ensure the full astrocyte was captured. Location of astrocyte within the corpus callosum was confirmed by the distinct linear arrangement of DAPI+ nuclei. Astrocytes outside of the corpus callosum were not imaged. Incomplete astrocytes were excluded from analysis.

Imaris 10.2 (Bitplane) software was used to trace the branching architecture of each astrocyte. Tracing was accomplished using the AI filament tracer function as described previously (Coble et al., 2026) with modifications for WM astrocytes. This machine learning approach was iteratively trained on a pilot dataset of eight images of individual astrocytes in flat mounted corpora callosi before implementing it for this analysis. Concentric Sholl spheres were placed every 5 µm around the traced astrocyte’s branches using a MATLAB based Sholl analysis Xtension for IMARIS (Gastinger, Oxford Instruments, 2021). CSV files containing Sholl intersection data were exported for each cell individually. This data was manually compiled into a single csv file and formatted for compatibility with a previously published In-Vivo-Sholl-Analysis R-script, available at https://github.com/Eroglu-Lab/In-vivo-Sholl-Analysis. Data were analyzed in Rstudio (version 2026.01.2+418) using a linear mixed-effects model to account for repeated measurements obtained at increasing radial distances from the soma. Experimental condition was included as a fixed effect, and repeated observations within cells were modeled as correlated measurements. Statistical significance of condition effects was assessed by ANOVA with Tukey’s post test of the fitted model, with p < 0.05 considered statistically significant. For visualization, mean Sholl profiles were plotted as the average number of intersections at each radius for each condition. Variability was represented using shaded ribbons corresponding to the standard error of the mean (SEM) at each radius.

### Endfoot Analysis

Endfeet were assessed in 100 μm-thick coronal sections containing corpus callosum, which were prepared for the 3D morphology analysis. IHC details and image acquisition were performed as described in that section. The presence of astrocytic endfeet was classified as either present or absent at each developmental timepoint in IMARIS 10.2 by visual assessment of 1) astrocyte process structure and 2) colocalization between the astrocyte processes and AQP4. Minimal post processing of the AQP4 channel (normalize layers) was applied to visualize AQP4 in the deeper planes of each image. Astrocytes lacking the AQP4 marker and an endfoot structure and were classified as “absent”. Astrocytes with a visible endfoot structure or colocalization between AQP4 and the astrocyte process were classified as “present”. We analyzed 39 images from 6 mice at P8, 30 images from 6 mice at P14, 36 images from 7 mice at P21 and 23 images from 6 mice at P70. To determine whether the proportion of endfoot-positive observations differed across developmental stages, a chi-square test of independence was performed using contingency tables of endfoot presence versus timepoint in Rstudio (version 2026.01.2+418). Standardized residuals were calculated for each cell of the contingency table to identify specific timepoints that contributed significantly to the overall association. Residuals with an absolute value greater than 1.96 were considered to indicate a substantial deviation from the expected frequency under the null hypothesis of independence. Data visualization was performed using stacked proportional bar graphs displaying the percentage of endfoot-present and endfoot-absent observations at each timepoint.

### Astrocyte Tiling Analysis

Tiling analysis was performed in 100 μm-thick coronal sections containing motor and somatosensory cortex (GM) and corpus callosum (WM). Tissue from CD1 mice previously intracortical injected with mCherry-CAAX AAV and GFP-CAAX AAV was collected at P14 and P21. IHC was performed using chicken anti-GFP primary antibody (1:1000, Fisher, GFP1010) and rabbit anti-RFP primary antibody (1:2000, Thermo Fisher, 600-401-379), along with goat anti-chicken Alexa Fluor 488 secondary antibody (1:200, Thermo Scientific, A-11039) and goat anti-rabbit Alexa Fluor 594 secondary antibody (1:200, Thermo Scientific, A11037). High magnification confocal z-stack images (1024 x 1024) containing whole astrocytes were acquired on an Olympus FV3000 microscope with a 30x objective and 2x optical zoom with a 0.5um step size. Criteria for acquisition required two neighboring cells labeled in opposite colors and in contact with one another, with at least one of these cells being fully contained within the tissue section. Images were excluded from analysis if 1) neither cell was whole 2) region of overlap was not included in the image 3) more than one cell body was identified in either astrocyte 4) one or both astrocytes was too dim for faithful territory reconstruction.

Imaris 10.2 (Bitplane) software was used as described above to calculate astrocyte territory volume (Baldwin et al., 2021; Eaker & Baldwin, 2022). Once a Convex Hull was constructed around both astrocytes, the Object-Object statistics were enabled for both convex hulls and the overlapping volume between the two convex hulls was exported directly from IMARIS. The total volume of each cell’s territory was also exported from IMARIS. These measurements were compiled in Excel for each image, and the overlapping volume was divided by the volume of the largest or the whole astrocyte to calculate the percent overlap between the two adjacent astrocytes. Percent overlap measurements were then arranged into a longform csv file for statistical analysis in Rstudio (version 2026.01.2+418). Values were only excluded from statistical analysis if the IMARIS constructed surface or convex hull did not faithfully represent the astrocyte’s shape or territory. In Rstudio, cleaned data was fit to a mixed-effects model with region as a fixed effect, animal as a random effect and overlap between cells as individual observations. This model was used to account for repeated measures between regions and variability in the number of observations per animal per region. Statistical significance of regional effects was assessed by ANOVA of the fitted model with Tukey’s post test, with p < 0.05 considered statistically significant. For visualization, individual cell values and animal averages were plotted in Prism 11.

### Node of Ranvier Density and Astrocyte-Node Interaction Analysis

Node of Ranvier density and Astrocyte-Node interactions were analyzed in 40 µm coronal sections containing corpus callosum. Tissue was collected from Aldh1l1-GFP mice at P14, P21, and P60. IHC was performed using chicken anti-GFP primary antibody (1:1000, Fisher, GFP1010), mouse IgG1 anti-CASPR primary antibody (1:100, DSHB, K65/35), and mouse IgG2a anti-AnkG primary antibody (1:100, DSHB, N106/36), followed by goat anti-chicken Alexa Fluor 488 secondary antibody (1:200, Thermo Scientific, A-11039), goat anti-mouse IgG1 Alexa Fluor 594 secondary antibody (1:200, Thermo Scientific, A-21125) and goat anti-IgG2a Alexa Fluor 647 secondary antibody (1:200, Thermo Scientific, A-21242). To enable analysis of astrocyte contacts, guinea pig anti-GLAST primary antibody (1:1000, Millipore Sigma, AB1783) and anti-guinea pig Alexa Fluor 488 secondary antibody (1:200, Thermo Scientific, A-11073) were added to the staining protocol for P60 tissue. High magnification confocal z-stack images (1024 x 1024) were acquired on an Olympus FV3000 microscope with a 60x objective and 2x optical zoom. Each image included 50 slices with a 0.43um step size.

Eight images were taken systematically for each of the four animals per timepoint. Two images were taken from four sections; one image was taken at the midline, and one to the right or to the left of the midline. Images were deconvolved using the constrained iterative deconvolution process in CellSens (Evident) software. The deconvolved images were converted to the IMARIS format and analyzed in IMARIS 11. Images were excluded when a software issue prevented conversion to the IMARIS format after deconvolution. Of the 96 images taken, 16 failed to convert. This error occurred at all timepoints, and erroneous files were excluded from the study resulting in a final data set of 3-8 images per animal from 4 animals at each timepoint: 24 total images at P14 cohort, 26 total images at P21, and 30 total images at P70. The analysis workflow in IMARIS 11 was customized using machine learning to train the software to identify and count Nodes of Ranvier (NOR) within defined regions of interest (ROI). Three regions of interest were hand drawn to encompass astrocytes processes within axon bundles for each image. In defining ROIs, astrocyte cell bodies were avoided in favor of regions with astrocyte processes and axon bundles, as identified by staining for NORs. To count NOR, a surface was constructed around the CASPR channel and spots were formed from AnkG channel. The AnkG spots were filtered first by quality to exclude all spots with an intensity less than 500 at its center, and then by proximity to the CASPR surfaces (Supplemental Fig. 4). NOR are between 1-2 µm long (Arancibia-Cárcamo et al., n.d.) with the concentration of AnkG appearing between the two CASPR rich regions flanking the node. To eliminate AnkG spots that were too far from CASPR to be NOR, all spots more than 1 µm away from the CASPR surfaces were discarded (Supplemental Fig. 4). Finally, a machine learning filter, which was iteratively trained on a pilot dataset of 24 images to recognize the dash-dot-dash pattern characteristic of NOR stained by Casper and AnkG, was applied to the AnkG spots to remove those that were not associated with CASPR in the correct pattern (Supplemental Fig. 4). To count astrocyte-node interactions, a surface was constructed around the astrocyte processes in each ROI. AnkG spots within 0.425 µm, twice the image’s resolution limit as calculated by the Rayleigh’s criterion, of an astrocyte’s process were counted as astrocyte-NOR contacts. The total number of NOR and astrocyte-NOR contacts from all three ROIs were divided by the total ROI volume to calculate densities per image for each measurement, and an interaction fraction was calculated by dividing the number of astrocyte-NOR contacts by the number of NORs per image. These values were input into Prism 11 for statistical analysis. Normality was assessed using the Shapiro–Wilk test and visual inspection of QQ plots. Statistical significance of changes in NOR density and astrocyte-NOR contact density were assessed by nested one way ANOVA with Sidak’s post test to compare timepoints. Statistical significance of the interaction fraction was assessed by nested t-test, with p < 0.05 considered statistically significant. Graphs were generated in Prism11 and show values from individual images and animal averages.

### Corpus Callosum Astrocyte Coverage Analysis

Astrocytic coverage of the corpus callosum was assessed in 40 µm coronal sections from Aldh1l1-GFP mice at P8, P14, P21, and P60. IHC was performed as described above using chicken anti-GFP primary antibody (1:1000, Fisher, GFP1010) and rabbit anti-GFAP primary antibody (1:1000, Agilent Technologies, Z033429-2), followed by goat anti-chicken Alexa Fluor 488 secondary antibody (1:200, Thermo Scientific, A-11039), goat anti-rabbit Alexa Fluor 594 secondary antibody (1:200, Thermo Scientific, A11037) and DAPI. Confocal images with 20 tiles were acquired with a 20x objective on the Zeiss LSM980 microscope to capture the thick region of corpus callosum around the brains’ midline. Images were acquired as a 10 µm z-stack with a 1 µm step size and multi-point z plane maps were used to adjust the center of each z-stack for unevenness or tilt in the sample. Three images were acquired from three animals per time point.

Images were analyzed using a custom Python-based macro in FIJI (version 2.61.0/1.54p). For each image, the DAPI channel was used to manually define a region of interest (ROI) around the corpus callosum. All measurements were restricted to the pixels within the ROI. Astrocyte density was determined by identifying nuclei positive for both DAPI and Aldh1L1-GFP. The nuclear channel was segmented using automated thresholding with the Otsu method followed by watershed separation of adjacent nuclei. The Aldh1L1-GFP channel was segmented used automated thresholding with the Mean method. Binary masks generated from the DAPI and Aldh1L1 channels were combined using the AND operation to identify colocalized nuclei. Colocalized nuclei were counted using particle analysis and density was calculated as the number of colocalized nuclei divided by the ROI’s area. The process coverage was quantified from the GFAP channel which was segmented using automated thresholding with the Huang method to generate a binary mask. The Process coverage fraction was calculated as the area occupied by GFAP positive pixels divided by the ROI’s area. To normalize the process coverage to the astrocyte density (cell per µm^2^), a coverage index (µm^2^ per cell) equal to the process coverage fraction divided by the cell density was calculated. Higher coverage index indicated greater process coverage per cell whereas lower values indicate reduced process coverage relative to cell density. Quality control outputs including thresholded masks for each channel, binary masks of colocalized nuclei and overlay images shows segmented objects on the corresponding raw fluorescence channels were generated for each image. These outputs were visually inspected to verify the analysis prior to downstream statistical analysis. Statistical analysis of density, process fraction and coverage index was performed in Prism 11. Normality was assessed using visual inspection of QQ plots and statistical significance was assessed by nested one way ANOVA with Sidak’s post test to compare timepoints, with p < 0.05 considered statistically significant. Graphs were generated in Prism11 and show values from individual images and animal averages.

### Synapse Imaging and Analysis

IHC to detect excitatory synapses was performed as described previously (Eaker et al., 2026), with modifications. 40 µm-thick coronal sections containing the somatosensory cortex and corpus callosum from Aldh1L1-eGFP mice at P14, P21, and P60 were used for analysis. IHC was performed using guinea pig anti-VGlut1 primary antibody (1:1000, Synaptic Systems 135 304), rabbit anti-PSD95 primary antibody (1:300, Thermo Fisher 51-6900), and chicken anti-GFP primary antibody (1:1000, Aves GFP-1020). To enable successful PSD95 labeling, Triton ampules were used with 2 weeks of opening to prepare TBST. The following secondary antibodies were used: goat anti-guinea pig IgG Alexa Fluor 647 (1:200, Thermo Fisher Scientific A-21240), goat anti-rabbit IgG Alexa Fluor 594 (1:200, Thermo Fisher Scientific A-11037), and goat anti-chicken IgY Alexa Fluor 488 (1:200, Thermo Fisher Scientific A-11039). High magnification confocal z-stack images were acquired using an Olympus FV3000 (60x objective, 1.64x optical zoom) inverted confocal microscope. For each animal, 3 z-stack images (15 slices, 0.34 µm step size) were acquired from layer 5 of somatosensory cortex and from the corpus callosum. Each image converted in 5 separate maximum projection images (MPI) of 3 slices each for a total of 15 MPIs per region per animal. Co-localized presynaptic and postsynaptic puncta were quantified using Synbot (Savage et al., 2024) with the following parameters: 2-channel, noise reduction, manual thresholding, minimum pixel = 3, pixel overlap. Images were collected from 3 animals per time point. Animal averages were analyzed using Prism 11. Normality was assessed using visual inspection of QQ plots and statistical significance was assessed by one way ANOVA with Sidak’s post test to compare tissue types within each timepoint, with p < 0.05 considered statistically significant. Graphs were generated in Prism11 and show values from animal averages.

## Quantification and Statistical Analysis

Statistical analyses were performed in GraphPad Prism 11 or Rstudio depending on the experiment. For each experiment, the number of subjects and specific statistical analyses are described in the methods above and in the figure legends. Sample sizes were determined based on previous or pilot experiments for each experiment. Specific details for inclusion and exclusion are included in specific methods subsections.

## Funding Statement

The Baldwin Lab is supported by the NIH, DP2 NS136873 to K.T.B. and T32GM133364 to K.H.M.H. The UNC Neuroscience Microscopy Core is supported in part by funding from the NIH-NICHD Intellectual and Developmental Disabilities Research Center Support Grant P50 HD103573. The BRAIN Initiative Viral Vector Core is supported in part by the NIH U24NS124025 to K. Ritola.

## Ethics Statement

All experimental protocols were performed in accordance with NIH guidelines and received approval from the Animal Care and Use Committee of UNC Chapel Hill.

## Data Availability Statement

Custom analysis files and scripts are available https://github.com/BaldwinLabUNC/Astrocyte_morphology. Raw data files available upon request.

## Conflict of Interest Statement

The authors declare no conflict of interest.

## Notes

### Competing Interest Statement

The authors have declared no competing interest.

