## Supplemental Figures 1-5 for "Morphological Hallmarks of White Matter Astrocytes Across Development"

| Comparison | Territory Volume | Surface Area | Relative Complexity |
|---|---|---|---|
| P8 GM vs. P14 GM | 0.0137 | 0.0005 | 0.0097 |
| P8 GM vs P21 GM | 0.0093 | 0.0107 | <0.0001 |
| P14 GM vs. P21 GM | 0.8921 | 0.2660 | 0.0153 |
| P21 GM vs. P70 GM | 0.2699 | 0.0005 | 0.0104 |
| P8 WM vs. P14 WM | 0.9030 | 0.0564 | <0.0001 |
| P14 WM vs. P21 WM | 0.1550 | 0.1832 | 0.3007 |
| P21 WM vs. P70 WM | <0.0001 | <0.0001 | 0.0552 |
| P8 GM vs. P8 WM | 0.9589 | 0.0136 | <0.0001 |
| P14 GM vs. P14 WM | 0.0127 | 0.0001 | 0.0009 |
| P21 GM vs. P21 WM | 0.1684 | 0.0574 | <0.0001 |
| P70 GM vs. P70 WM | 0.0027 | 0.6580 | <0.0001 |

**Supplemental Table 1:** ***Uncorrected p-values for morphology metrics comparing GM and WM astrocytes at P8, P14, P21 and P70***

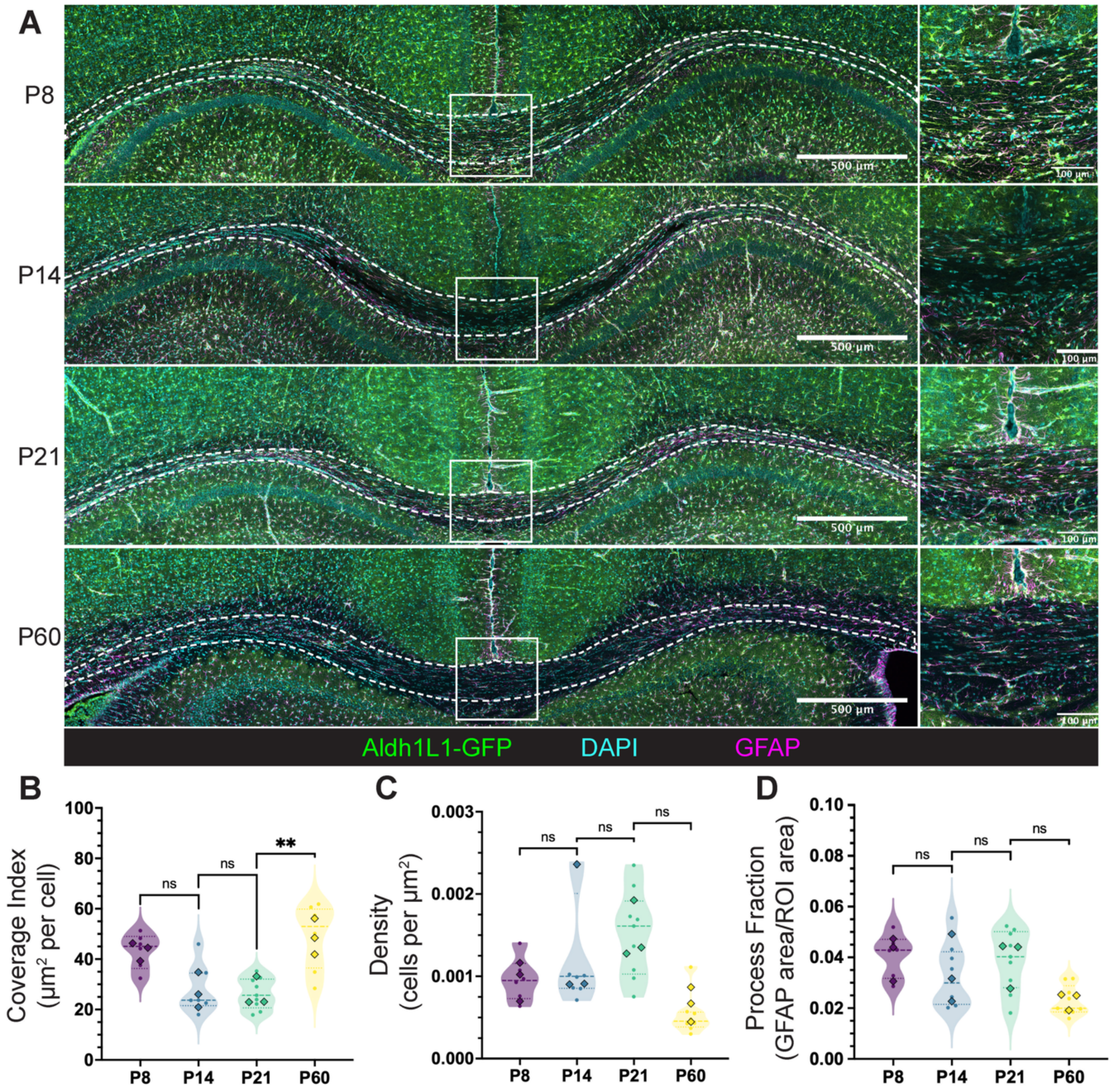


**Supplemental Figure 1:** ***Astrocyte coverage of the corpus callosum increases into adulthood***

**A)** Representative tiled confocal images of astrocytes dual labeled with Aldh1L1-GFP (green) and GFAP (magenta) and nuclei (DAPI, cyan) in the corpus callosum (outlined by dotted white lines) at P8, P14, P21 and P60. Higher magnification regions focused on the midline of the corpus callosum are shown to the right and are highlighted on the full images with white boxes. **B-D)** Violin plots of **B)** coverage index **C)** astrocyte density and **D)** process fraction in the corpus callosum. Dots represent individual images and diamonds represent per animal averages. 2-3 images were analyzed from 3 animals per timepoint. Nested one way ANOVA with Sidak's post test. ns = p<0.05, * = p≤0.05, ** = p≤0.01, ***= p≤0.001, **** = p≤0.0001.

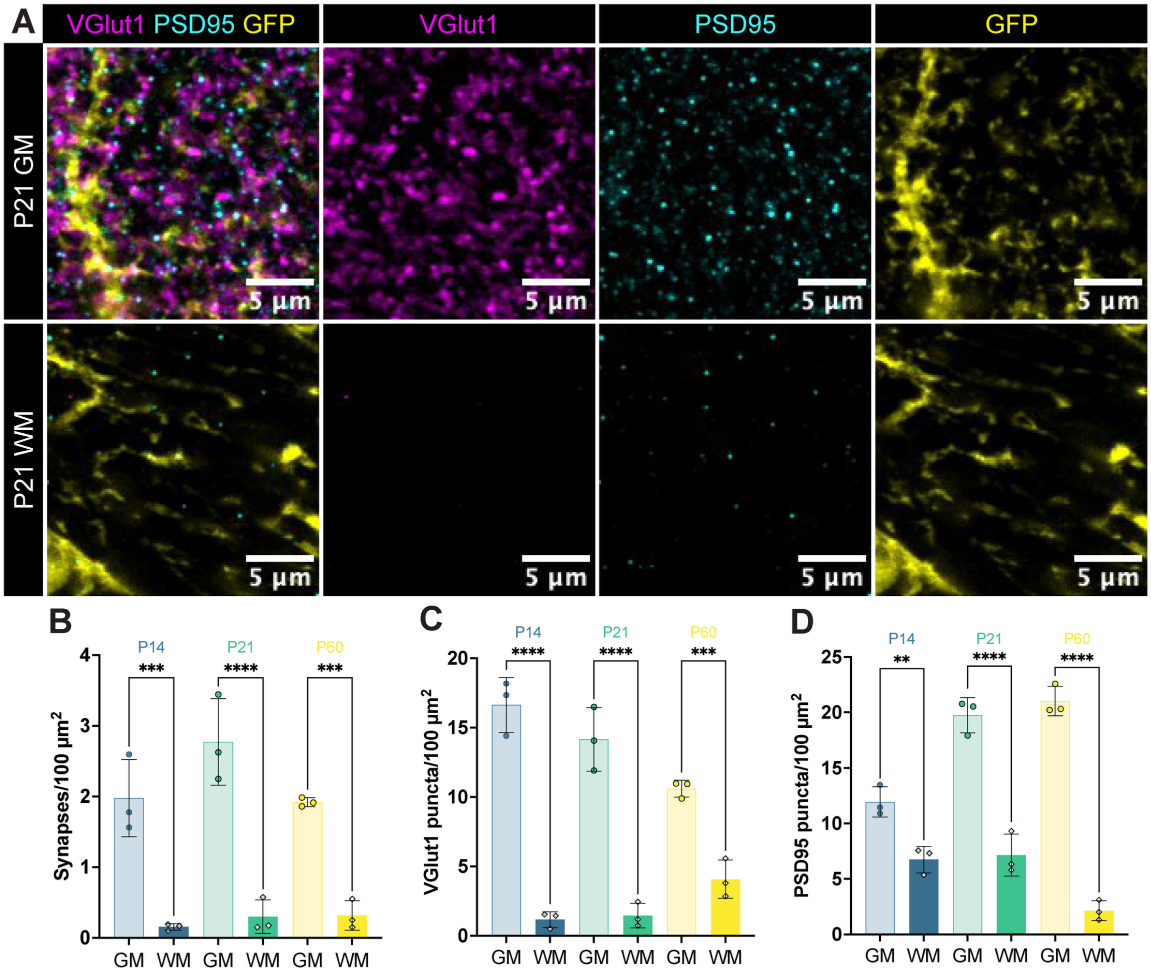


**Supplemental Figure 2:** ***The corpus callosum is largely devoid of synapses***

**A)** Representative confocal images of presynaptic marker VGLUT1 (magenta), postsynaptic marker PSD95 (cyan) and astrocytes (Aldh1L1-eGFP, yellow) in the layer 5 somatosensory cortex and corpus callosum at P21. **B-D)** Bar graphs showing the density of **B)** co-localized VGlut1 and PS95 (synapses/100 µm$^2$) **C)** VGlut1 and **D)** PSD95 in the cortex and corpus callosum at P14, P21 and P60. Diamonds represent per animal averages, error bars show the ±SD. 3 images from 3 animals per timepoint. One way ANOVA with Sidak's post test. ns = p<0.05, * = p≤0.05, ** = p≤0.01, ***= p≤0.001, **** = p≤0.0001.

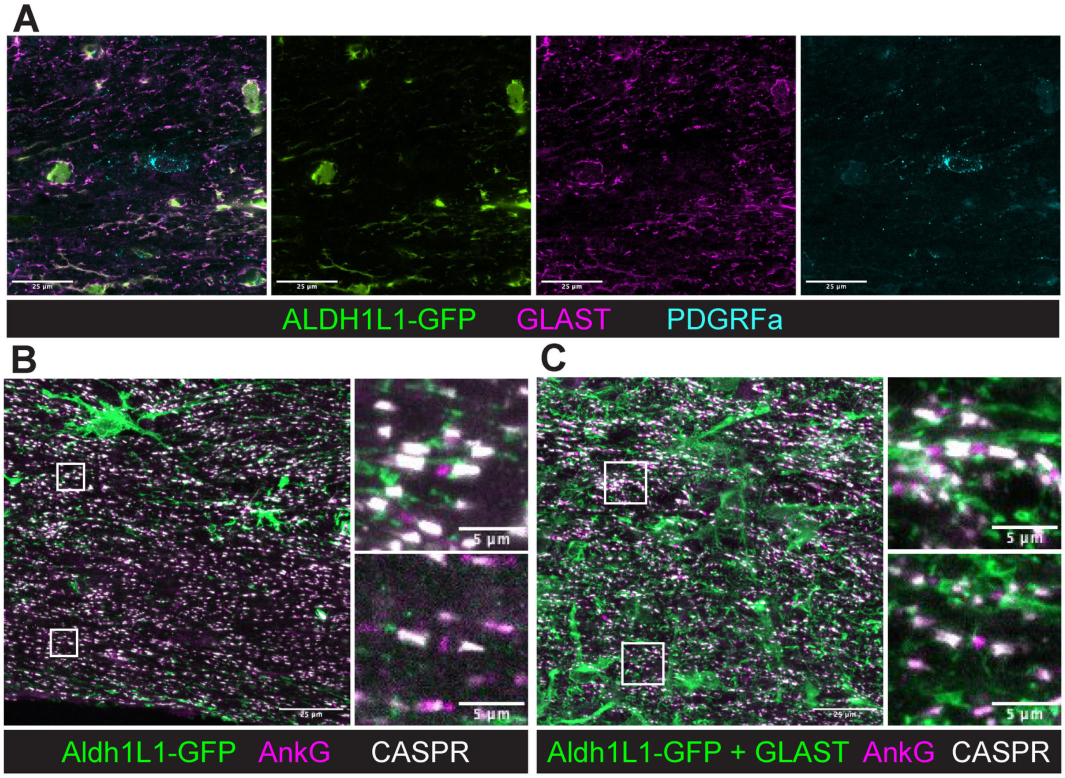


**Supplemental Figure 3:** ***GLAST effectively labels fine WM astrocytes processes into adulthood without OPC colocalization***

**A)** Representative image of staining for Aldh1L-eGFP (green), GLAST (magenta) and PDGRF$\alpha$ (cyan) at P60 in the corpus callosum. **B-C)** Representative images of **B)** Aldh1L1-eGFP labeled and **C)** Aldh1L1-eGFP + GLAST labeled astrocytes and NOR (visualized by AnkG and CASPR) at P60. Higer magnification insets, highlighted with white boxes on the full images, show examples of astrocyte-NOR interactions with main branches (top) and fine branches (bottom).

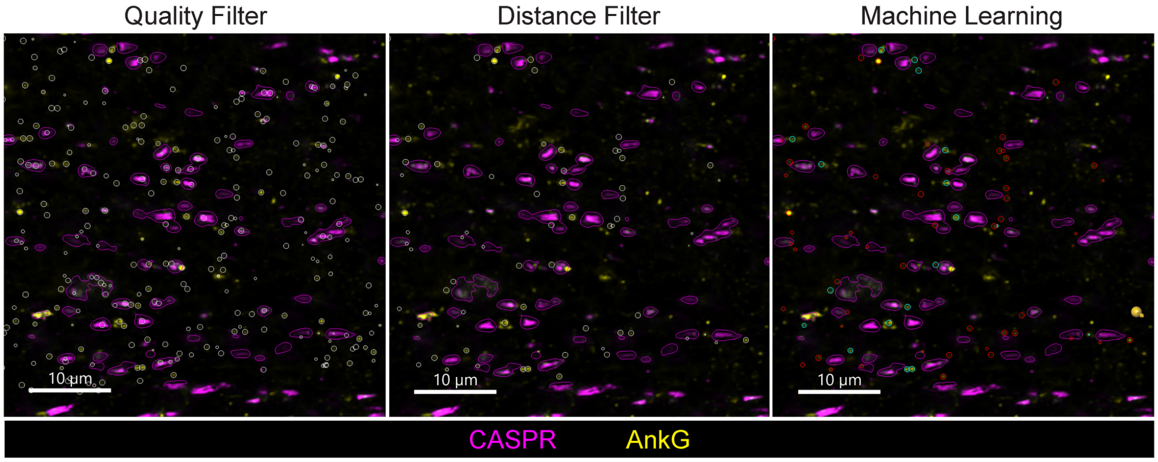


**Supplemental Figure 4:** ***An IMARIS-based machine learning filter identifies Nodes of Ranvier in vivo.***

Imaris 11 based workflow for filtering spots generated from the AnkG channel to specifically mark Nodes of Ranvier. Spots were filtered first by quality, then by their distance to surfaces generated from the CASPR channel, and then by a machine learning-based filter trained to recognize the dash-dot-dash pattern of CASPR and AnkG staining. The red dots are discarded by the filter while the blue dots are kept and counted.

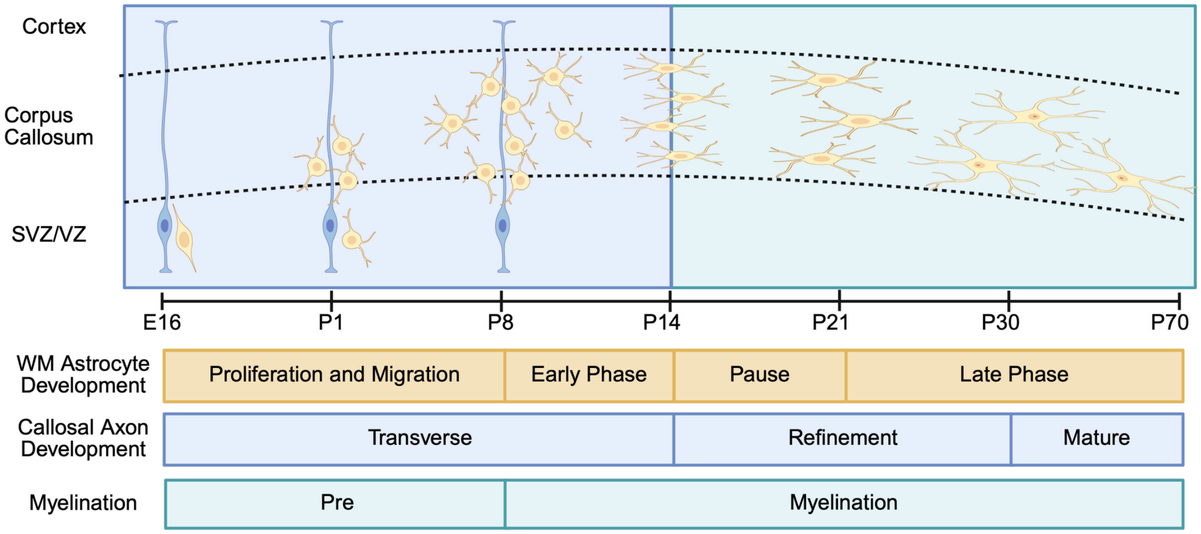


**Supplemental Figure 5:** ***A biphasic model for the development of astrocytes in the corpus callosum*** Created with BioRender.com.
